# Symmetric Inheritance of Histones H3 in Drosophila Male Germline Stem Cell Divisions

**DOI:** 10.1101/2021.10.22.465494

**Authors:** Julie Ray, Keith A. Maggert

## Abstract

Mitotically-stable “epigenetic” memory requires a mechanism for the maintenance of gene-regulatory information through the cell division cycle. Typically DNA-protein contacts are disrupted by DNA replication, but in some cases locus-specific association between DNA and overlying histones may appear to be maintained, providing a plausible mechanism for the transmission of histone-associated gene-regulatory information to daughter cells. Male *Drosophila melanogaster* testis germ stem cell divisions seem a clear example of such inheritance, as previously chromatin-bound Histone H3.2 proteins (presumably with their post-translational modifications intact) have been reported to be retained in the germ stem cell nuclei, while newly synthesized histones are incorporated into daughter spermatogonial chromosomes. To investigate the rate of errors in this selective partitioning that may lead to defects in the epigenetic identity of germ stem cells, we employed a photoswitchable *Dendra2* moiety as a C-terminal fusion on Histones H3 (Histone H3.2 and Histone H3.3); we could thereby discriminate histones translated before photoswitching and those translated after. We found instead that male germ line stem cell divisions show no evidence of asymmetric histone partitioning, even after a single division, and thus no evidence for locus-specific retention of either Histone H3.2 or Histone H3.3. We considered alternative hypotheses for the appearance of asymmetry and find that previous reports of asymmetric histone distribution in male germ stem cells can be satisfactorily explained by asynchrony between subsequent sister stem cell and spermatogonial divisions.

## RESULTS AND DISCUSSION

Epigenetic mechanisms are conceptualized to be divided into establishment and maintenance phases (1,2). During the former, inducers and/or transcription factors effect changes in chromatin structure, including chemical modifications to nucleosomal histones at specific chromosomal loci, altering expression of the underlying DNA. During the latter, those established chromatin structures are imagined to be retained through DNA replication, allowing local gene regulatory statuses to persist through the cell division cycle and leading to long-term mitotically-stable (and/or meiotically-transmitted, transgenerational) inheritance even in the absence of the establishing effector. It is a key feature of epigenetic inheritance that even identical sequences can act disparately because the regulatory information is indifferent to the underlying sequence (3) and different epigenetic information (*e*.*g*., activating or repressing transcription, conferring centromeric activity) can associate with individual chromosome loci. However these ideas are not without ongoing controversy (3, 4). The essence of epigenetic inheritance – and thus the focus of the controversy – is the maintenance phase, especially elucidating if and how epigenetic information can survive the unpacking, replication, and repackaging of chromatin during S-phase.

Epigenetic gene regulation has been proposed to assure the stability of cell identity, with instability leading to dedifferentiation, precocious or aberrant determination, loss of pluripotency, transdetermination, or the onset of disease states (5). Perhaps by analogy to the immortal strand hypothesis (6, 7), some have envisioned the evolution of a system in which histone-mediated epigenetic information is preferentially retained in stem cells during cell division, safeguarding their pluripotency and limiting errors in maintenance that might occur during multiple rounds of “copying” histone-encoded epigenetic information (8).

Reports of such systems in *Drosophila* germ stem cells of the testes (9) and somatic stem cells of the gut (communicated in (10)) provide experimentally manipulable opportunities to investigate how epigenetic and stem cell division asymmetries relate. Male germ stem cells were reported to retain older Histone H3.2 (*i*.*e*., those that had been chromatin-bound during the preceding G1-phase), while the first non-stem daughter/sister cell, the primary spermatogonial “gonialblast,” received newly transcribed and translated Histone H3.2 during S-phase. Tran and colleagues suggested that this asymmetry explains how “epigenetic information could be maintained by stem cells or reset in their sibling cells that undergo cellular differentiation” (9) and Zion and Chen reiterated that this could “… directly link the asymmetric histone inheritance mode with the establishment of distinct cell identities after one cell division”(10). Our interest in transgenerational epigenetic inheritance (*i*.*e*., from one organismal generation to the next) drew our attention to the former study because of the opportunity to specifically erase epigenetic information through mitotic exchange (11-12), and to correlate defects in non-random histone segregation in the germ line stem cell with epigenetic instabilities in offspring (13-15). For example, Tran and colleagues reported that many cell-pairs contained some degree of both old and new histones. These observations led us to wonder whether these cells might represent natural epigenetic instabilities, and we could potentially identify sperm developmental lineages that were prefigured to exhibit epigenetic instability in the next organismal generation (as in (15)).

To address this possibility, we obtained flies expressing either *histone H3*.*2*::*dendra2* or *histone H3*.*3*::*dendra2* fusion genes from Dr. Amanda Amodeo (16). Expression of both were under control of their natural genetic regulatory elements (Figure 1A) and were kept as homozygotes to provide two expressing copies per genome. This system has advantage over the heat shock-inducible and tissue-specific GAL4-controlled expression of monochromatically-labeled H3 histones used in other studies (Figure 2A) because expression of either H3::Dendra2 in the germ line should be relatively free from artifacts related to over-expression, ectopic expression (*i*.*e*., outside of the natural cell-type, outside of the natural expression phase in the cell-cycle – S-phase for Histone H3.2 and G1-G2 for Histone H3.3 (17)), chimeric 5’ and 3’ control sequences including the 3’ stem-loop (18), or perdurance of Histone H3, GAL4, or FLP mRNAs or proteins from previous cell generations. Such artifacts may be considerably disruptive, for example in yeast over-expression of histones, or buildup of nucleoplasmic histones by mutation of *spt16*, leads to a downregulation of *CLN3* and G1 delay (19-20), and in *Drosophila* over-expression can disrupt proper cell cycle timing in embryonic divisions (21) or to intergenic suppression (17). With respect to histone function, the Amodeo laboratory reported that mitosis occurs with expected timing and duration in early embryogenesis (16, 21), and therefore concluded that mitotic functions are not significantly stalled or impaired by the fluorescent moieties. In our experience, flies expressing these chimeric proteins do not express any viability, fertility, or morphological defects, and develop in concert with heterozygous siblings and *wild*-*type* strains.

**Figure 1.**
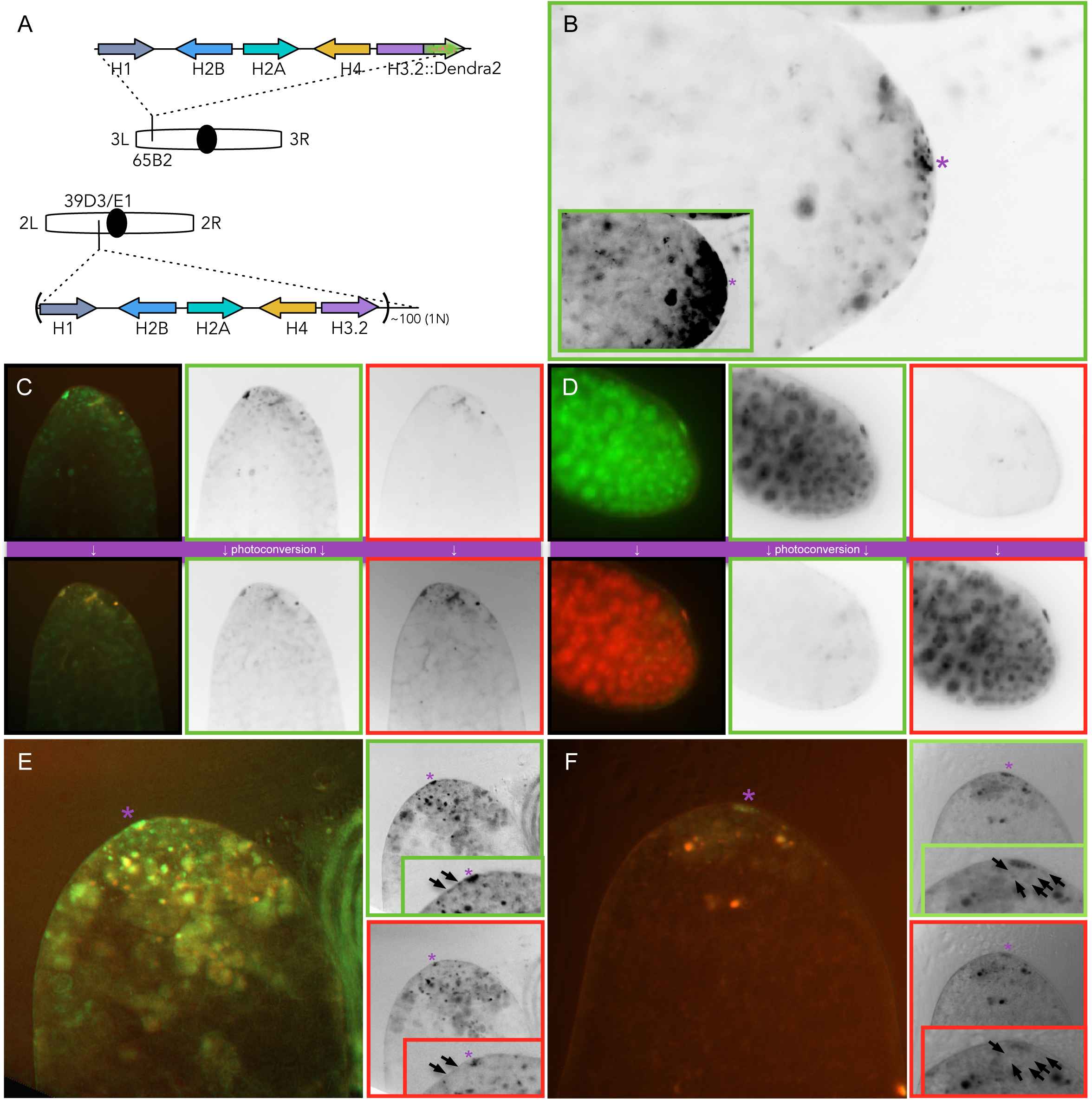
Photoswitchable fusion proteins in live tissue allow high temporal resolution in studying histone dynamics. **(A)** Diagram of transgene containing one copy of the histone gene complex, including Histone H3.2 fused at its C-terminus to Dendra2. In this case, the transgene is incorporated via *PhiC31 attP*-mediated integration at cytological band 65B2, and uses its natural regulatory elements for expression. The endogenous *histone* locus at 39D3/E1 has approximately 200 copies (as a diploid). **(B)** Pronounced expression of Histone H3.2::Dendra2 fusion in non-germinal hub cells (asterisk), low levels in germ line stem cells and primary spermatogonia, and a lower level in the subterminal testis where spermatogonial cysts are found. Image is presented as linear inverted monochrome and adjusted bright and contrast (inset) to enhance visibility. **(C)** Top row of images show fluorescence from Histone H3.2::Dendra2 prior to photoconversion, and inverted monochromatic channel separations (green fluorescence in the middle column, and red in the right column). Bottom row shows the same testis and separations after photoconversion. **(D)** As in (C), but showing expression and photoconversion of a Histone H3.3::Dendra2 fusion. **(E)** Testis from male expressing Histone H3.3::Dendra2 fusion, dissected, photoconverted, then aged 40 hours before imaging. Linear inverted monochrome separations for green (top) and red (bottom) are shown. Insets show magnified view of stem and 1° spermatogonial cells (arrows). **(F)** As in (E) except testes were aged 20 hours after dissection and photoconversion, before imaging.

**Figure 2.**
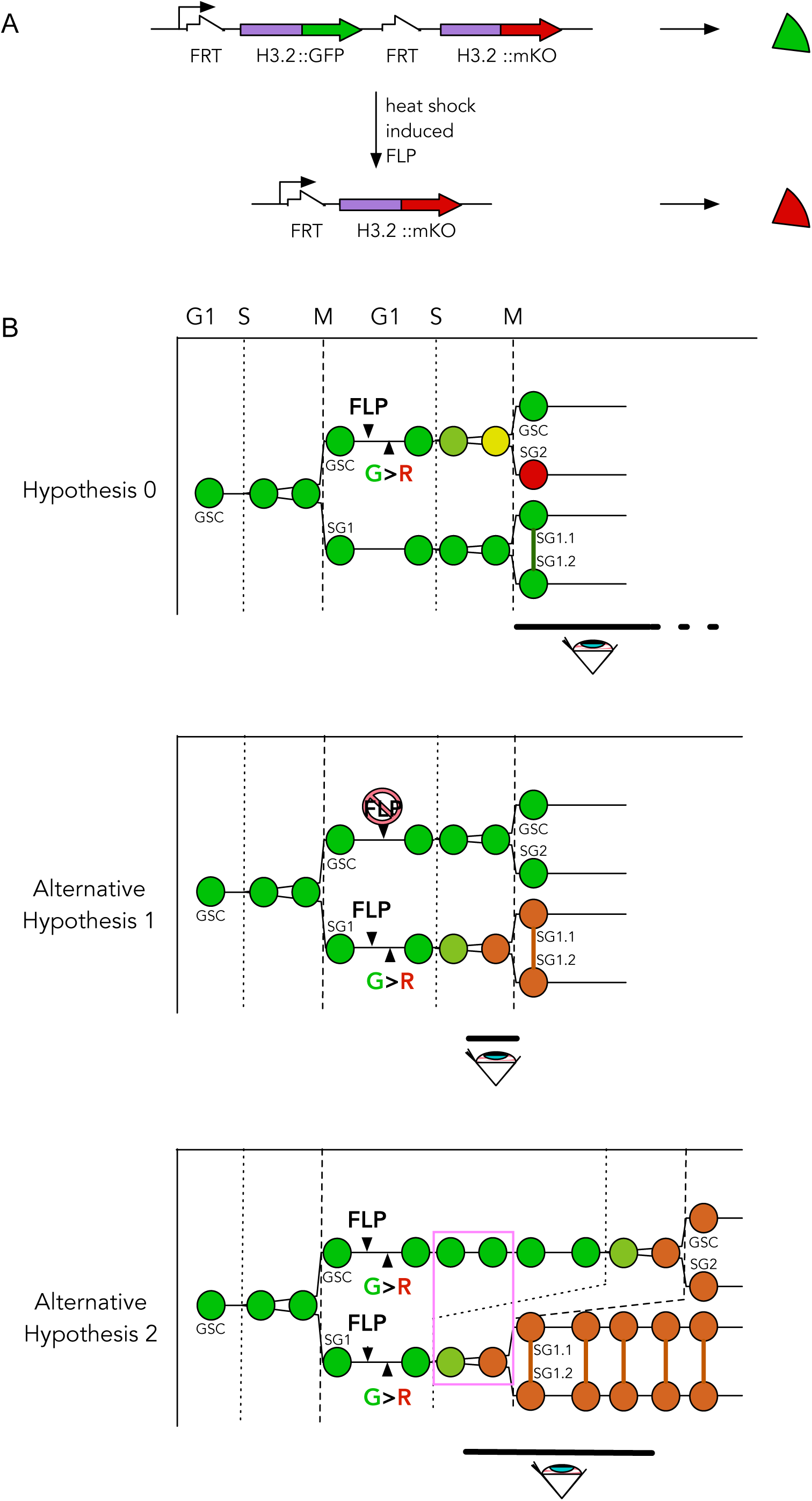
Alternative hypotheses to explain the appearance of histone retention on “old” chromosomes in *Drosophila* male germ cells. **(A)** The gene constructs used in previous studies, in which UAS-linked labeled H3.2::GFP was replaced by a labeled H3.2::mKO by a FLP-dependent genome rearrangement; expression was controlled by tissue-specific GAL4 expression in the germ line (8). **(B)** The germ line stem cell (GSC) and primary spermatogonium (SG) (*a*.*k*.*a*., “gonialblast”) nuclei are colored according to the scheme in (A): nuclei/chromosomes with “old” histones (having been used previously to package chromatin in G1-phase (G1), and may retain their post-translational modifications, in particular those that are gene-regulatory) are green and “new” histones (newly-translated and naïve in regard to histone modification) are red. <u>Hypothesis 0</u>, the model proposed by Chen and colleagues (8, 9, 24, 26), in which older histones are specifically retained during S-phase (S) on the chromatids fated to remain in the germ line stem cell after mitosis (M). After heat-shock-controlled FLP expression (“FLP”) and FLP-dependent genome rearrangement (“G>R”), green fluorescent histones represent the older histones and are retained in the germ line stem cells, whereas the newly-expressed red fluorescent histones are partitioned to the spermatogonium (SG1). The vertical line connecting spermatogonia represent incomplete cytokinesis of the developing cyst; such incomplete cytokineses continue to ultimately produce 16-nuclei cysts during spermatogenesis. The black-bar-and-eye indicate the time of observation when the two nuclei (GSC and SG) are distinct. Two alternative hypotheses can account for the appearance of asymmetric histone inheritance. <u>Alternative Hypothesis 1</u>, the FLP-induced genome rearrangement to replace green-labeled histones with red-labeled histones may be inefficient (or absent, slashed “FLP”) in germ line stem cells (GSC), while it proceeds (more) efficiently in spermatogonia (SG1). Cell-limited rearrangement produce germ line stem cells and sister spermatogonia after S-phase expressing discordant fluorescence. The frequency of such cell pairs would be a function of the difference in FLP induction efficiencies in each. Note also that this hypothesis, which does not involve exclusive partitioning of old and new histones to old and new chromatids, predicts some green fluorescence (in addition to the dominant red) in spermatogonia if some of the old histones are reused in S-phase (indicated by orange color in SG1.1 and SG1.2). <u>Alternative Hypothesis 2</u>, the S/G2-phases subsequent to the one that first separates the stem/spermatogonial sister and spermatogonial cysts (*i*.*e*., the GSC<–>SG2 and SG1.1<–>SG1.2) are temporally offset. This could be a difference in the time of entry into S-Phase (as shown) or a different rate of S-Phase (faster in SG1 than in GSC). Discordancy in fluorescence is indicated as a pink rectangle. This hypothesis also predicts some green fluorescence in spermatogonial nuclei (denoted by the orange color in SG1.1 and SG1.2). Note that <u>both</u> alternative hypotheses need not be mutually exclusive. But one clear point of departure from Hypothesis 0 from the alternatives is the retention of fluorescent asymmetry in the germ line stem cells after protracted time (in hypothesis 0) *versus* eventual symmetry (in the alternatives).

We observed that the fluorescence of the Histone H3.2 fusion protein was reliably detectable in the germline stem cells but only at very low levels (Figure 1B-C). While expression was detected in the germinal stem cells and primary spermatogonia, decreased gradually as spermatogonial cysts developed, and was undetectable by the 16-nuclei cyst stage in our experiments. Since histones are oftentimes re-used between subsequent S/G2 phases, we consider the possibility that the low-and-diminishing levels of fluorescence through pre-meiotic spermatogenesis reveals a protein dilution from the site of highest gene expression, the germline stem cells. Low expression in the stem cells is likely owed to the relatively high multiplicity (about 100-fold) gene copy number of endogenous *histone H3*.*2* genes in the *histone* gene complexes (22). However, it is also possible that Histone H3.2 is not the predominant histone in these cells, cells instead relying on H3.3 expression which was expressed robustly and uniformly in germinal spermatogonia, germinal cysts, and non-germinal cyst cells (Figure 1D). This is consistent with the discovery that H3.3 mutants express a meiotic phenotype in *Drosophila* males (17). In support of this possibility, *histone H3*.*2*::*dendra2* was robustly expressed in the small hub cells adjacent to the stem cells (Figure 1B, asterisk), indicating that fluorescence of the single copy of *histone H3*.*2*::*dendra2* is not intrinsically limited by competition amongst the gene expression of unlabeled endogenous *histone H3*.*2*. We photoconverted dissected *histone H3*-*Dendra2*-expressing testes and confirmed photoconversion by low-resolution imaging of whole testes, and by close inspection of a subset of mounted testes (Figure 1C-D). Testes from *histone H3*.*2*-*dendra2*-expressing males were set in culture medium for 40 hours after photoconversion. At the end of that *in vitro* incubation period, whole live testes were inspected at both the red- and green-emitting wavelengths, allowing us to analyze live tissue and use the fluorescence of the Dendra2 moiety to discriminate those histones derived from pre-S-phase chromatin (red) from those that were synthesized *de novo* and incorporated during subsequent S-phases (green).

Germ line stem cells and spermatogonia were identified by location in the apex of the testis and their juxtaposition to the cluster of hub cells (23). Of the 24 whole testes we inspected after photoconversion and culturing, in no case did we find red fluorescent Histone H3.2 limited to germ line stem cells (Figure 1E). Instead, we observed *de novo* (green fluorescent) Histone H3.2 exclusively in the stem cells and paired spermatogonia. We could confirm that the lack of red-fluorescence was not due to inadequate photoconversion by the presence of red fluorescence in the non-mitotic hub cells (asterisk), and also by the presence of red fluorescence in secondary spermatogonia and cysts. The latter further indicates that photoconversion, dissection, and culture *ex vivo* did not terminally arrest cell division. Thus we could find no evidence for preferential retention of previously-used Histone H3.2 molecules in germ stem cells. We repeated the experiment with a shorter (20-hour) post-photoconversion culture time and as before we could detect no evidence for retained histones in the germ stem cells (Figure 1F). It appeared that all cells examined had completely replaced red-fluorescent Histone H3.2 with *de novo* (green-fluorescent) Histone H3.2, within the detection limits of our assay system. Some germline stem cells had a low level of red fluorescence detectable along with green fluorescence, indicating either incomplete photoconversion, or photoconversion during S-phase (Figure 1H).

In general, photoconverted red Histone H3.2-containing nuclei were visible in many cells, indicating that these histones and their photoconverted fluorescent moieties, are not intrinsically unstable. As a control, we performed similar photoconversion-chase experiments using the *Histone H3*.*3-Dendra2* fusion gene. As expected, after 40 hours in culture the red fluorescence had disappeared from the germline stem cells and spermatogonia. All of the nuclei that contained photoconverted Histone H3.3 also had newly-translated Histone H3.3, indicating active processes of replacement either by transcription or replication. This experiment is consistent with the work of Tran and colleagues, and further supports the longevity of the photoconverted Dendra2 moiety.

We considered the disparity in ours and Tran and colleagues’ findings, considering their experimental system and how it differed from ours. In their experiments, heat-shock was used to induce a ubiquitous expression of the FLP site-specific recombinase, excising the GFP-labeled Histone, and activating the tissue-specific expression of a mKO-labeled Histone (Figure 2A). We reasoned two alternative hypotheses for the appearance of histone retention reported in their work (Figure 2B, “Hypothesis 0”). First, we considered that the heat shock inducible FLP recombinase used to replace the labeled Histone H3.2 may be less efficient in stem cells than in spermatogonia. In this way, it seemed feasible that switching might occur in spermatogonia (conferring them red fluorescence) and not in the germ stem cells (where green fluorescence would be retained) (Figure 2B, “Alternative Hypothesis 1”). Second, a difference in the timing of cell division between germ stem cells and primary spermatogonial daughter cells might create the appearance of histone retention if the spermatogonia underwent a subsequent S-phase (and thus histone incorporation) prior to the germ stem cell’s next S-phase (Figures 2B, “Alternative Hypothesis 2”).

To test the first alternative hypothesis, we generated males bearing the same heat shock inducible FLP used by Tran and colleagues (9), as well as the G-TRACE lineage marking system (24). G-TRACE uses a FRT-mediated chromosome rearrangement to excise an intervening STOP cassette, irreversibly activating a ubiquitously-expressed GFP upon exposure to FLP; the genome rearrangement itself, excision of an extrachromosomal circle, is quite similar to the one performed by Tran and colleagues (Figure 2A). We heat-shocked third instar larvae and confirmed that all cells of later-metamorphosed adult testes were fluorescent green, indicating that FLP was efficiently induced in all germ cells in the primordial gonad (Figure 3A). Notably, the germ line produced only green fluorescent stem cells, spermatogonia, spermatocytes, spermatids, and sperm, indicating that larval male gonadal stem cells efficiently expressed FLP under heat shock control. Next, we heat-shocked intact adults for 1 hour and analyzed them for fluorescence 72 hours later. If FLP was efficiently expressed in adult germ stem cells, we expected entirely green fluorescent cysts derived from germ stem cell divisions after heat shock. If FLP was inefficiently expressed, we instead expected to see no or incompletely-penetrant expression in germ stem cells and recently derived cysts depending on whether secondary spermatogonia or spermatocytes efficiently express FLP and GFP. We observed the first outcome and could find no small cysts or germ stem cells without GFP expression (Figure 3B), indicating efficient expression of FLP and subsequent genome rearrangement in the germ stem cells. To rule out the possibility that FLP-induced expression of GFP was preferentially in cysts, but heat-shocked testes experienced developmental arrest at that stage, we limited FLP expression by reducing the time of heat shock to 15 minutes. After 72 hours we observed stochastic GFP activation in a subset of cysts (Figure 3C), indicating definitively that we induced FLP-mediated GFP activation in the germline without otherwise affecting expression in clonal descents. This also allowed us to unambiguously identify the number of nuclei in expanding cysts, showing that FLP and GFP were definitively within in the germline rather than the encasing cyst cells; in fact we counted only 2^n^ cells in these labeled cysts, indicating that genome rearrangement did not occur in the non-germinal cyst cells. Collectively, we interpret these data as refutation of our first alternative hypothesis, and confirmation that the appearance of asymmetric histone inheritance cannot be explained by unequal heat shock inducible FLP activity in germ stem cells and primary spermatogonia.

**Figure 3.**
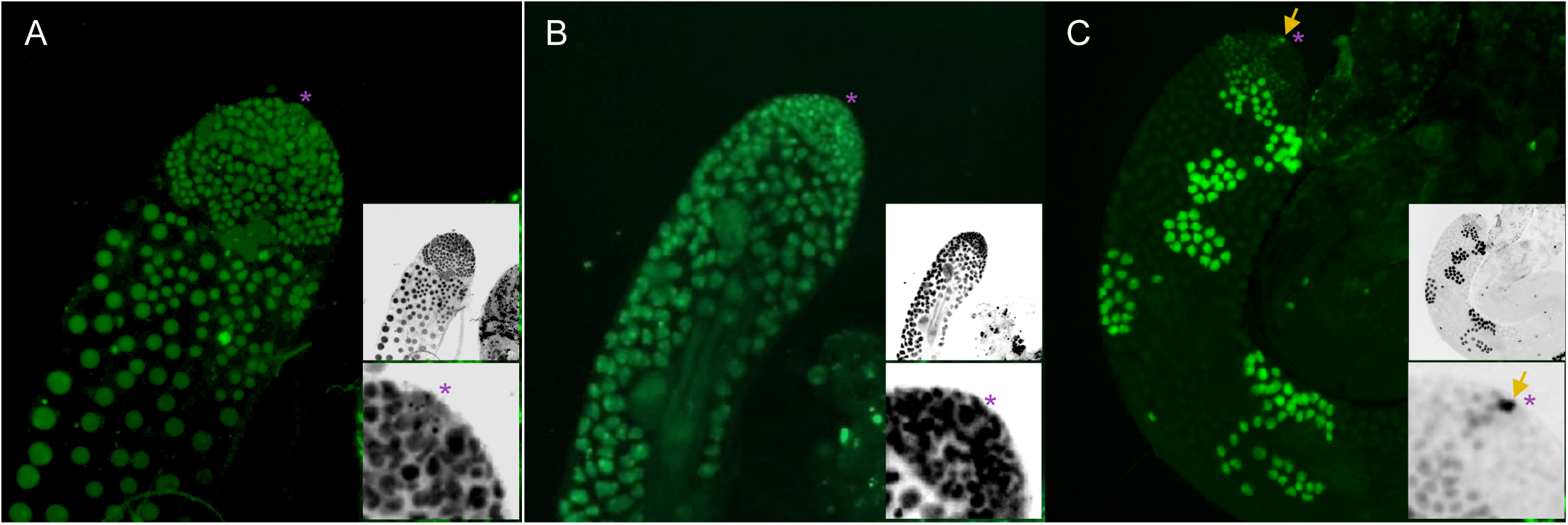
Heat-shock induction of GAL4 transactivator and FLP-mediated genome rearrangements are efficient in germ line stem cells. **(A)** Male testes containing a hsp70-GAL4 heat shock inducible GAL4 gene and the G-TRACE lineage tracer, heat shocked as a third instar larvae then dissected as adults 3 days after eclosion. Green fluorescent cells indicate genome rearrangement and permanent ubiquitous activation of the GFP gene in all primordial germ cells. **(B)** Male testes of the identical genotype heat shocked for 2 hours as an adult and dissected 120 hours later, also indicating all germ line stem cells efficiently underwent FLP-dependent genome rearrangement. **(C)** As in (B) but heat shocked for 10 minutes, limiting FLP expression. In this case only one germ stem cell (arrow) expressed sufficient FLP to induce the permanent activation of ubiquitously-expressed GFP. In all panels, insets are inverted images of the same testis and higher-magnification view with hub cells marked with asterisks.

We note that it is not possible with our experimental design to rule out a subset of this alternative hypothesis, that FLP-mediated excision occurs in S- or G2-phase of the cell cycle, and that one sister chromatid experiences the FLP-mediated excision and the other does not. Such an occurrence could also manifest as an asymmetry, like that observed by Tran and colleagues. However, in that case, we would expect one half of the exchanged chromosomes to be segregated to the stem cell and one half to the sister spermatogonium. We found no such corroborating evidence in 72-hour post-heat shock testes. Further, we are aware of no study that describes such limitations for FLP-mediated recombination, and in fact FLP-mediated excision seems very efficient even in G1 somatic cells and of meiotic secondary spermatocytes (11); on these grounds we reject this alternative hypothesis.

To test the second alternative hypothesis (Figure 2B), we analyzed cell cycle timing in germ stem cells and primary spermatogonia using two live cell cycle reporters. First, we looked for asynchrony in the onset of S-phase of germ stem cells and their paired spermatogonia using a PCNA-GFP fusion reporter line. We gently squashed testis tips to better reveal the amount and localization of PCNA-GFP, and found that expression of nuclear PCNA was varied in the testes tips, indicating considerable heterogeneity in the phases of the cell cycle. Specifically, nuclear PCNA-GFP-positive spermatogonia could be found adjacent to non-fluorescent hub-adjacent stem cells (Figure 4A-C). We interpret this to mean that spermatogonial cells may enter S-phase (and thus active Histone H3.2 incorporation) independent of the time at which the sister stem cell does. This observation and interpretation accords with analyses by Matunis and colleagues, which showed that germline stem cells and primary spermatogonia are no longer linked at the time the latter undergoes cell division (25). Because male secondary spermatogonia incompletely divide in every cell division after the germline stem cell division from which they arise, ultimately forming cysts of 2^n^ cells, up to 16 interconnected spermatogonia (Figure 4C), we interpreted any individual cells near the tip of the testes that are not in contact with the small hub cells to be primary spermatogonia.

**Figure 4.**
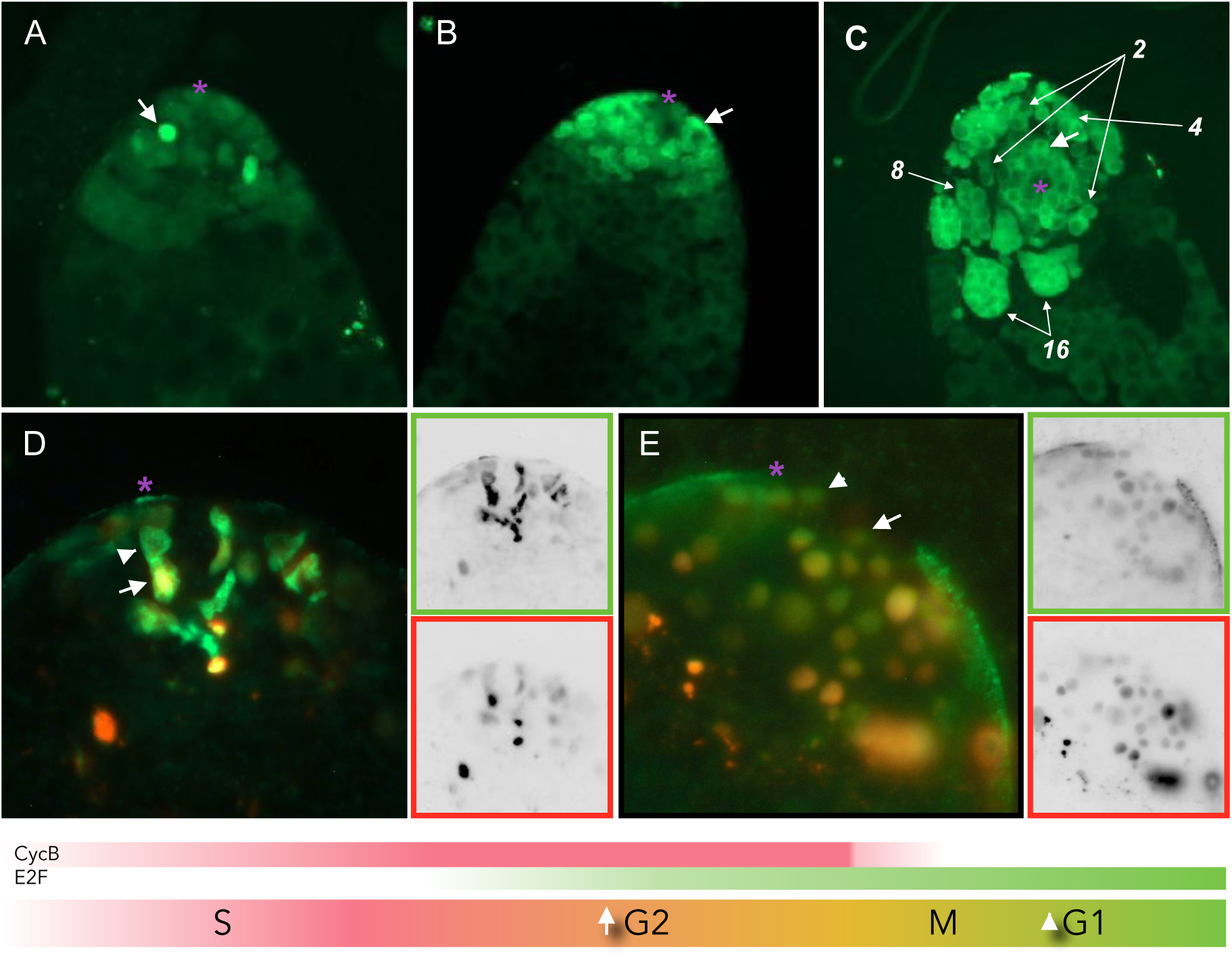
Germ line stem cells and sister primary spermatogonia have discordant subsequent cell cycles. **(A), (B), (C)** Testes from males bearing a homozygous transgene encoding a PCNA::GFP fusion, dissected 3 days after eclosion. Nuclear fluorescence indicates a cell in S-Phase. Asterisks mark hub cells, and arrows indicate spermatogonia in S-Phase whose sister germ line stem cells (adjacent to both arrowhead-indicated and asterisk-indicated cells) are not. These are denoted in Figure 2B (“Alternative Hypothesis 2,” pink rectangle) **(D), (E)** Testes from males bearing a germ line stem cell specific GAL4 transgene and the UAS-FlyFUCCI cell division cycle reporter, dissected 3-5 days after eclosion. In this system, RFP is fused to a Cyclin B degradation signal and GFP to an E2F degradation signal (diagram below (D) and (E)). Cells in S-Phase fluoresce red, those in G2 fluoresce red-to-yellow, and those in mitosis or G1 fluoresce green. Insets show monochrome separations (green on top, red below) adjusted for brightness to show fluorescence. White arrowheads indicate stem cells in M/G1-Phase and white arrows indicate sister spermatogonia in S/G2-Phase. Numbers indicate spermatogonial cysts with indicated number of synchronous spermatogonia.

Second, we employed tissue-specific Fly-FUCCI (26) to monitor stages of the cell cycle for cells in the germ line. This reporter consists of a GAL4-dependent GFP moiety fused to an E2F-derived degradation signal and an RFP moiety fused to the degradation signal from Cyclin B, such that cells in G1 fluoresce green, cells in S-phase fluoresce red, and cells in G2 or mitosis appear as the overlap (Figure 4D). Expression was limited to the germline by use of a germ-specific GAL4 transgene characterized elsewhere (27). We dissected testes from males three days after eclosion and analyzed the germ stem cell niche and niche-adjacent cells for cell cycle phase. We could observe pairs of cells wherein the hub-proximal cell was in G1 (or early S) and the latter was in G2 (Figures 4E-F). We cannot ascertain whether these asynchronies are in S-phase entry or the rate of S-phase – it does not matter in terms of critically testing Alternative Hypothesis 2 as these conditions would appear identical – however we favor the latter since G1 cells are uncommon in germ stem cells.

These two methods to monitor the cell divisions of germ stem cells and sister primary spermatogonia lead us to the same conclusion, that there is independence of cell cycle timing between the germ stem cell’s and the primary spermatogonium’s next divisions. Our observations establish that asynchrony in cell division does occur in male testis. We expect from first principles that asynchrony in S-phase initiation or rate in paired spermatogonia and sister germ stem cell would lead to asymmetry in replication-coupled incorporation of Histone H3.2. Further, we interpret that in the case of Tran and colleague’s work, even though sister germ stem cells and spermatogonia may have rearranged genomes and carry the *H3*.*2-mKO* reporter, because Histone H3.2 is limited to replication-dependent incorporation (23, 28) only those cells that undergo the an S-phase will appear red. This is not true of the replication-independent incorporation of Histone H3.3, and indeed both they and we see even incorporation of “old” and “new” Histone H3.3 in all germinal cell types of the testes. We expect that alterations to cell cycle timing may affect the asynchrony between stem cell and daughter spermatocytes, giving the appearance of disrupted asymmetric inheritance (29).

It has been challenging to envision a model whereby histones “know” which are the chromatids destined for the spermatogonium and which will stay with the germ stem cell. Any model would require that within each replication bubble, new histones must remain associated with leading strand synthesis at one fork and lagging strand synthesis at the other. But as every daughter chromatid is a product of alternating leading- and lagging-strand synthesis of the nascent strand, it must also be that all of the origins along the chromosome conspire to pick which strand is destined for the spermatogonium versus germ stem cell, and those coordinations must be agreed-upon by every fork on a chromosome, and by every chromatid in the nucleus. Recent data demonstrating biased histone incorporation at forks appear orders of magnitude lower than would be necessary to explain even asymmetry, what more complete histone partitioning (30-31), nor do those analyses reveal any coordination between origins. *Post facto* repair of “old and new” information (as in (32)) is ruled out by both cytological and genetic experiments. Yadlapalli and colleagues showed that chromosome strands are not preferentially segregated during the same cell division (male germ stem cells) (33), arguing against preferential labeling of “immortal strands” by some chemical mark. This is consistent with work by Dan Lindsley and colleagues showing that chemically-induced single-strand lesions are not preferentially retained in germ stem cells (34).

Finally, we offer two final challenges to asymmetric histone inheritance in male germ stem cells. First, asymmetric retention has no apparent function as it has long been established that depleted germ stem cells can repopulate the germ stem cell niche by recruiting and “de-differentiating” primary spermatogonia by re-establishing hub-dependent JAK/STAT signaling (36, 37). This is a normal and frequent process occurring at detectable frequencies in wild-type testes, yet there is no indication that these cells (or the organisms derived from their fated sperm) express any phenotype. Second, Tran and colleagues report an asymmetry of about 80%, yet even an asymmetry of 90% (that is, 90% of dividing stem cells show complete retention of “old” histones), it would take a mere 7 cell divisions to be effectively random [(0.9)^5^ = 0.59; (0.9)^6^ = 0.53; (0.9)^7^ = 0.49]; at 80%, it would take 3 divisions to erase any benefit of asymmetric histone inheritance. Each germ stem cell in *Drosophila* undergoes approximately 70 divisions in the reproductive lifetime of the organism (1.5 divisions per day over an approximately 45 day reproductive lifespan), and probably at least 5-6 before a male can find a mate in the wild.

We conclude that there is no evidence of asymmetric or preferential histone incorporation in *Drosophila* testis germ stem cells, nor have any phenotypes been described that would indicate or take adaptive advantage of such a system. This challenges the notion that “epigenetic” differences exist between the germ stem cells and primary spermatogonia, and likely the wealth of literature describing the role of the niche in determining the germ stem cell fate is sufficient to explain the maintenance of these cells.

## MATERIALS AND METHODS

### Dissections, Culturing, and Imaging

Dissection of flies was performed between one and five days after eclosion in Ringer’s solution or Phosphate-Buffered Saline (PBS) after a brief 70% ethanol wash to remove cuticular organics. Testes were imaged in PBS as whole-mounts or gently-squashed samples and imaged on a Zeiss AxioZoom.v16 using Filterset 38HE (Excitation 470/40, Splitter 485, Emission 525/50) for green fluorescence and Filterset 00 (Excitation 530-585, Splitter 600, Emission 600-) for red fluorescence. Photoswitchable Dendra2 moieties were photoconverted with a 60-second exposure to 380-400 nm light (Filterset 49) on a Zeiss AxioSkopII.mot at low magnification (5X). Dissected and photoconverted testes were cultured in testis culture medium supplemented with penicillin, streptomycin, and insulin. Most Images were adjusted for brightness but not contrast to retain linearity of signal, expect where low-level fluorescence was highlighted as indicated in figure legends. JPEGs were exported and processed in Adobe Photoshop to create inverted monochrome separations.

### *Drosophila* Strains

The H3.2 and H3.3 fusion strains were a gift from Amanda Amodeo. The heat shock inducible GAL4 was P{ry[+t7.2]=hsFLP}1, y[1] w[1118]; +; Dr[1]/TM3, Sb[1]; +. G-TRACE was w[*]; +; P{w[+mC]=UAS-RedStinger}6, P{w[+mC]=UAS-FLP.Exel}3, P{w[+mC]=Ubi-p63E(FRT.STOP)Stinger}15F2; +. PCNA::GFP was w[1]; +; P{w[+mC]=PCNA-EmGFP}T137; +. Germ line stem cell GAL4 was w[1118]; +; P{w[+mC]=GAL4::VP16-nos.UTR}CG6325[MVD1]; +. UAS-FlyFUCCI was w[1118]; Kr[If-1]/CyO, P{ry[+t7.2]=en1}wg[en11]; P{w[+mC]=UASp-GFP.E2f1.1-230}64 P{w[+mC]=UASp-mRFP1.NLS.CycB.1-266}5/TM6B, Tb[1]; +.

## ACKNOWLEDGEMENTS

We gratefully acknowledge support from the National Institutes of Health to the laboratory (R01GM123640), the University of Arizona Cancer Center (P30CA023074), and the Bloomington Drosophila Stock Center (P40OD018537), as well as the University of Indiana Univeristy to the last. We also acknowledge outstanding resource support from the University of Arizona (for the University of Arizona Genetics Core) and the Univeristy of Arizona Cancer Center in the form of core services. Finally, to Dr. Amanda A. Amodeo for the Histone-Dendra2 fusion strains.

## Notes

### Competing Interest Statement

The authors have declared no competing interest.

### Summary of Updates

Some new images, some new words.

